# Face inversion reveals holistic processing of peripheral faces

**DOI:** 10.1101/112433

**Authors:** Petra Kovács, Balázs Knakker, Petra Hermann, Gyula Kovács, Zoltán Vidnyánszky

## Abstract

Face perception is accomplished by face-selective neural processes, involving holistic processing that enables highly efficient integration of facial features into a whole face representation. It has been shown that in face-selective regions of the ventral temporal cortex, neural resources involved in holistic processing are primarily dedicated to the central portion of the visual field. These findings raise the intriguing possibility that holistic processing might be the privilege of centrally presented faces and could be strongly diminished in the case of peripheral faces. We addressed this question using the face inversion effect, a well established marker of holistic face processing. The behavioral results revealed impaired identity discrimination performance for inverted peripheral faces scaled according to the V1 magnification factor, compared to upright presented faces. The size of peripheral face inversion effect (FIE) was comparable to that found for centrally displayed faces. Face inversion affected the early ERP responses to faces in two time intervals. The earliest FIE was most pronounced in the time window between 130-140 ms following stimulus presentation, for both centrally and peripherally displayed faces and in the latter case, it was present only over the contralateral hemisphere. The timing of the next component FIE corresponded closely with the temporal interval of the N170 ERP component and showed strong right hemisphere lateralization, both when faces were displayed in the left or right visual field. Furthermore, we also showed that centrally presented face masks impaired peripheral face identity discrimination performance, but did not reduce the magnitude of the FIE. These findings revealed robust behavioral and neural inversion effects for peripheral faces and thus suggest that faces are processed holistically throughout the visual field.

**Highlights:** Robust behavioral and neural inversion effect was found for peripheral faces.

P1 ERP component is modulated by inverted central and contralateral faces.

N170 ERP component is modulated by centrally and peripherally presented faces.

Neural face inversion effect shows strong right hemisphere lateralization.

Faces are processed holistically throughout the visual field.

## 1. Introduction

Human face perception is fast and highly efficient. A growing body of experimental results suggests that it is subserved by a specialized network of cortical regions with face-selective neural responses, the face processing network (Calder & Young, 2005; Haxby, Hoffman, & Gobbini, 2000; Ishai, 2008; Ishai, Schmidt, & Boesiger, 2005). One hallmark of face perception that distinguishes it from the perception of other visual objects is holistic processing (Farah, Wilson, Drain, & Tanaka, 1998; Jacques & Rossion, 2009; Maurer, Grand, & Mondloch, 2002; Piepers & Robbins, 2012; Rossion, 2008). Holistic face processing refers to the processes enabling the perception of a face as an integrated whole. Although the extensive research in the last decades led to a remarkable progress in our understanding of behavioral and neural processes of holistic face perception (EEG: Caharel et al., 2013; Jacques & Rossion, 2009; Nemrodov et al., 2014, fMRI: James, Arcurio, & Gold, 2013; Zhang, Li, Song, & Liu, 2012; Zhao et al., 2014), there are several important issues that remained unexplored. In particular, since previous research on face perception focused on the processing of faces presented in the central visual field, the basic question whether and to what extent perception of peripherally located faces is accomplished by face-selective neural processes involving holistic processing remains unexplored. This question appears to be especially relevant in the light of recent research suggesting that visual object processing in the human ventral temporal cortex (VTC) is spatiotopically organized (Amedi, Malach, Hendler, Peled, & Zohary, 2001; Grill-Spector & Weiner, 2014; Hasson, Levy, Behrmann, Hendler, & Malach, 2002; Malach, Levy, & Hasson, 2002). Important to the findings of the current study, prior neuroimaging studies testing spatiotopy in the VTC found that the representation of objects whose recognition depends on the analysis of fine details – such as faces - is associated with a central visual-field bias. It has been shown that in the fusiform face area (FFA), the region of the VTC selectively involved in face perception, including holistic processing, nearly all neural resources are dedicated to the central (~7 deg) portion of the visual field (Kay, Weiner, & Grill-Spector, 2015). These findings raise the intriguing possibility that the highly efficient face-selective neural resources, which might be responsible for holistic face processing, might be the privilege of centrally presented faces and could be strongly diminished for peripheral faces.

To address this question, we tested the effect of picture-plane inversion on behavioral and EEG responses to both foveal and peripheral faces. The face inversion effect (FIE) is one of the most widely used markers of the highly developed face-specific visual processing skills in the case of foveal faces (Goffaux & Rossion, 2007; Rossion, 2008; Van Belle et al., 2010; Yin, 1969). It has been shown that inversion leads to slower and less accurate recognition and discrimination of faces by impairing holistic processing (Rossion, 2008, 2009; Rossion & Gauthier, 2002). In accordance with its behavioral effect, face inversion also affects the early components of the ERP responses, including the P1 as well as the N170 components, associated with structural processing of facial information (Bentin, Allison, Puce, Perez, & McCarthy, 1996; Colombatto & McCarthy, 2016; Eimer, 2000; Jacques, d’Arripe, & Rossion, 2007; Rossion, Gauthier, Tarr, & Despland, 2000; Rossion, Joyce, Cottrell, & Tarr, 2003). Both the P1 and the N170 ERP components emerge later with increased amplitudes for inverted relative to upright faces (Bentin et al., 1996; Eimer, 2000; Linkenkaer-Hansen et al., 1998; Rossion et al., 2000). This is thought to reflect the increased processing demands required for the integration of relational information among features within the face-selective cortical regions (Goffaux & Rossion, 2007; Rossion, 2008) and/or engagement of cortical regions not belonging to the core face network (for review see Yovel, 2016), such as the object-selective lateral occipital cortex and the parietal cortex. Importantly, the only study (McKone, 2004) we found in the literature that involved the picture-plane inversion of peripheral face stimuli obtained a strong and consistent FIE on face identification performance even at the largest tested stimulus eccentricity (21 deg), suggesting that processing of peripheral faces might be accomplished by face-selective neural mechanisms, also involving holistic processing.

In the present study we aimed at investigating the behavioral and electrophysiological effects of peripheral face inversion and comparing them to the foveal FIE. To this end, we measured ERP responses and eye movements while participants performed a three-alternative forced choice (3AFC) face identity discrimination task. During the task, faces appeared with either upright or inverted orientation in three different positions (in the center, and at an eccentricity of 10° in the left visual field (LVF) or right visual field (RVF)). Peripheral face stimuli were scaled according to the cortical magnification factor (Horton JC & Hoyt WF, 1991). The results revealed strong FIEs both on the behavioral and ERP responses to peripheral face stimuli. Additionally, we performed an additional behavioral experiment to test whether neural processes dedicated to the central visual field are involved in the face inversion effects obtained for peripheral face stimuli as well. This possibility has been raised by recent results (Fan, Wang, Shao, Kersten, & He, 2016; Williams et al., 2008) showing that peripheral object perception involves a foveal processing component mediated by feedback signals to the visual cortex representing the central visual field that can be revealed by presenting foveal noise shortly after the presentation of the peripheral target objects. We reasoned that if peripheral holistic face processing is mediated by neural processes dedicated to the central visual field, adding foveal masks, in addition to modulating the overall face discrimination performance, should strongly reduce the size of the peripheral FIE. Our results revealed that foveal upright and inverted faces, but not noise masks impair peripheral identity discrimination performance strongly for both upright and inverted faces. Importantly, however, the magnitude of the peripheral FIE was not affected by the foveal face masks. Taken together our findings provide support for the holistic processing of peripheral faces.

## 2. Methods

### 2.1. Participants

Eighteen healthy subjects participated in the experiment. Two participants were excluded due to inadequate fixation (see *Eye Tracking Data Acquisition and Analysis*), therefore 16 participants were included in the final analysis (6 male, 2 left-handed, mean ± SD age: 23.13 ± 3.4 years). All of them had normal or corrected-to-normal vision. None of them reported any history of neurological or psychiatric disease. They provided written informed consent in accordance with the protocols approved by the United Ethical Review Committee for Research in Psychology (EPKEB), Budapest, Hungary.

### 2.2. Stimuli

Twelve frontal images of male faces with neutral expression were generated using FaceGen Modeller 3.5 (Singular Inversions Inc., Toronto, Canada) and imported to Matlab 2008a (MathWorks, Natick, MA). Images were converted to grayscale and cropped to exclude external features such as hair or external contours and were presented through an oval aperture on a uniform, mid-grey background. The luminance and contrast of the images were equated with a custom-written Matlab script, adjusting the mean intensity values of the image to midgrey (grayscale intensity: 128). From the twelve faces, four different triplets were created, one for each of the four experimental blocks. We created 12 linear morph continua from the mean face of FaceGen to each of the identities. For morphing, a warp algorithm (Winmorph 3.01) was used. The morph level was set at 80% (where 0 means the average face and 100% is the original, unmorphed identity) based on a pilot study to yield an average accuracy of ~80-85% for upright faces presented at the center. Stimuli were presented in three different positions (center of the screen, and centered at an eccentricity of 10° (closest edge to center) in the left visual field (LVF) or right visual field (RVF)). Peripherally presented faces were scaled in size according to the cortical magnification factor (CMF, Horton JC & Hoyt WF, 1991) to compensate for the decrease in the cortical representation of the visual field towards periphery. The calculation of the image size was carried out with the formula M_linear_=A/(E+e_2_) described by Horton & Hoyt (1991) using A=29.2 mm, e_2_=3.67° as parameters calculated for human V1, where A is the cortical scaling factor measured in mm, E is the eccentricity in degrees, and e2 is the eccentricity (degrees) at which the stimulus subtends half the cortical distance that it subtends at the fovea (Dougherty et al., 2003; Horton & Hoyt, 1991). The image at the center subtended a 4 × 3 degrees of viewing angle, while images on the periphery were displayed according to the obtained CMF at the size of 14.9 × 11.17 degrees (Fig.1.).

**Fig.1.**
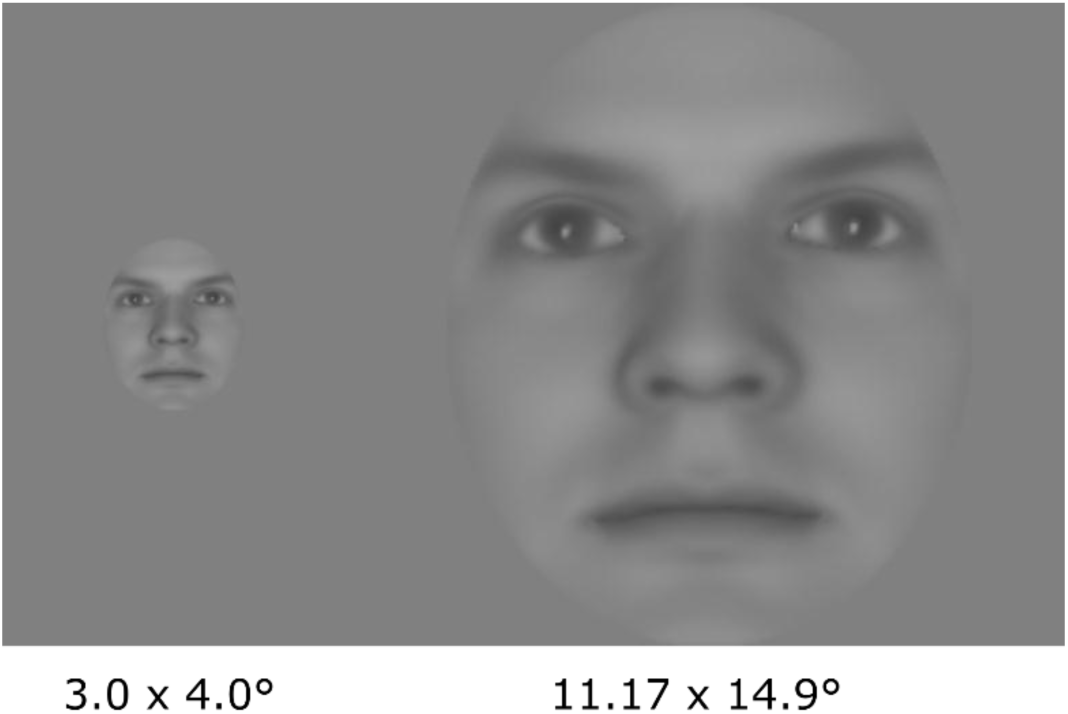
Example of the experimental face stimuli. Size of stimuli on the left was applied for centrally presented stimuli, while stimulus size on the right was used for peripheral stimuli. The size ratio of the stimuli is respected in the figure.

Participants viewed the stimuli on a 26 inch LCD monitor at a refresh rate of 60 Hz from 56 cm viewing distance, maintained using a chinrest. The experiment was controlled by Matlab (Mathworks, Natick, MA) using Psychtoolbox 3.0.9 (Brainard, 1997; Pelli, 1997).

### 2.3. Experimental Procedure

Participants were required to recognize the identity of the presented face in a three-alternative forced choice (3AFC) task. A fixation dot was present at the center of the screen during the entire experiment and participants were asked to fixate this dot regardless of where the stimulus appeared. In each experimental block, the 3AFC face discrimination task was performed on one of the four face identity triads. The triads were the same across all paticipants. The four identity triads were assigned to the first four blocks in random order, then in the remaining four blocks, they reoccurred in the same order.

In the training phase at the beginning of each block, participants learned the three identities from the face triad of the current block. First, the three unmorphed identities were shown side by side to allow participants to familiarize with them and to memorize their mapping to the three response keys (‘left arrow’, ‘bottom arrow’ or ‘right arrow’ on a QWERTY keyboard). After they felt confident with their knowledge, a subsequent practice phase started where the previously studied faces were presented individually, and participants were instructed to indicate face identity by pressing one of the three keys. Face stimuli disappeared after response, and were immediately followed by a feedback message displayed at the center of the screen (‘OK’, ‘MISSED’). Each of the three identities were repeated four times, and the learning phase was repeated when there was more than one erroneous response. After properly accomplishing this training phase, participants went on to the test phase.

In the test phase, after an intertrial interval of 1300-1700 ms, each trial started with a target face stimulus that was presented for 150 ms. Faces were shown upright or inverted randomly with equal probability. Participants were instructed to indicate face identity by pressing one of three keys as fast and accurately as possible. After stimulus offset they had 2000 ms to make their response. After their response, participants received feedback by the fixation point changing its color (green in the case of correct and red in the case of incorrect responses). The experiment consisted of 1152 trials, 192 trials per condition. Participants were allowed to take a short break between the blocks. The experiment lasted approximately one and a half hour.

### 2.4. Data Analysis

#### 2.4.1. Analysis of the Behavioral Data Collected During the EEG Acquisition

Responses and reaction times (RTs) were collected during the experiment and were submitted to a repeated-measures analysis of variance (ANOVA) with Orientation (Upright (UP), Inverted (INV)) and Position (Left Visual Field (LVF), Center (C), Right Visual Field (RVF)) as within-subject factors. The Greenhouse-Geisser correction was applied to correct for possible violations of sphericity. Post hoc tests were computed using Tukey’s honestly significant differences (HSD) procedure. Statistical analyses were carried out by STATISTICA 13.1 (StatSoft, Tulsa, OK).

To test the relationship between central and peripheral FIEs, we subtracted accuracy for inverted from that of upright faces for each position (LVF, C, RVF) separately. We conducted a correlation analysis between these difference indexes for presentation positions to test whether there is an association of behavioral FIE among different positions across participants.

For the correlation analyses we used the Robust Correlation Toolbox (Pernet, Wilcox, & Rousselet, 2013) and calculated skipped Pearson correlation coefficients. This method detects and removes bivariate outliers according to the adjusted box-plot rule (Pernet et al., 2013). The number of removed outliers are noted as ‘NO’ when reporting correlation results. 95% confidence intervals (CI) were calculated based on 10,000 resamples with the percentile bootstrap method implemented in the toolbox. To compare whether correlation coefficients differed between conditions we used Zou’s confidence interval (Zou, 2007) implemented in the *cocorr* software (http://comparingcorrelations.org; Diedenhofen & Musch, 2015).

#### 2.4.3. Electrophysiological Acquisition and Procedure

For the EEG data acquisition a BrainAmp Standard (BrainProducts GmbH, München, Germany) amplifier was used. Sixty-three active electrodes (ActiCap, BrainProducts GmbH, München, Germany) were placed according to the extended 10-20 international electrode system and mounted on an elastic cap with 1 additional periocular electrode under the right eye to record the electrooculogram. All channels were referenced to the TP10 electrode online and later digitally transformed to an average reference of all electrodes. The ground electrode was placed at the FPz electrode position. All impedances were kept below 5 kΩ. Data were sampled at 500 Hz. Data were acquired with BrainVision Recorder (version 1.2, BrainProducts).

#### 2.4.4. Electrophysiological Data Analysis

For pre-processing, EEGLAB 13.4.4.b (Delorme & Makeig, 2004) and ERPLAB 5.0 (Lopez-Calderon & Luck, 2014) were used running on Matlab R2015a (Mathworks, Natick, MA). A digital Butterworth zero-phase high-pass filter with half-amplitude cutoff frequency of 0.1 Hz and a slope of 12 dB/octave and a 24 dB/octave Butterworth zero-phase low-pass filter with a half amplitude cutoff frequency of 30 Hz were applied. Parks-McClellan 50 Hz notch filter was used to minimize the line-noise artefacts. All filtering wasdone using the ERPLAB toolbox.

The ERPs were calculated as follows. The continuous filtered EEG data were segmented into 800 ms epochs starting from 200 ms preceding stimulus onset for baseline correction. The average voltage measured during this pre-stimulus window was subtracted from the entire waveform. For artefact rejection we used a moving-window peak-to-peak threshold on 50 ms long windows with a voltage threshold of 100 μV. Following this automatic artefact detection, we performed artefact rejection by visual inspection as well. The cleaned data was averaged to obtain ERP waveforms for each subject and condition separately. For visualization, ERPs of the subjects were also averaged to compute grand averages.

Statistical analysis was performed on early peaks (P1, N170) of the average ERP waveform for each subject. The mean peak amplitudes and latencies were measured from waveforms averaged within electrode clusters. For the two investigated components the electrode clusters consisted of electrodes over both the right and left hemispheres, based on previous studies (Bankó, Gál, Körtvélyes, Kovács, & Vidnyánszky, 2011). Electrode clusters for the P1 component consisted of PO7, PO9, O1, and O9 for the left and PO8, PO10, O2, and O10 electrodes for the right hemisphere; for the N170 component, PO7, PO9, P7, and P9 was used over the left and PO8, PO10, P8, and P10 over the right hemisphere.

Amplitude values were analyzed separately for each component with three-way repeated-measures analysis of variance (ANOVA) with Hemisphere (Left Hemisphere (LH) and Right Hemisphere (RH)), Orientation (UP, INV), and Position (Ipsilateral (IL), Center (C), Contralateral (CL)) as within-subject factors. The levels of the position factor reflect the position of the stimulus relative to cerebral hemispheres: for the left hemisphere, LVF stimuli are coded as Ipsilateral and RVF stimuli are coded as Contralateral, and vice versa for the right hemisphere. This arrangement facilitates the interpretation of the data. The Greenhouse-Geisser correction was applied to correct for possible violations of sphericity. For post hoc tests Tukey’s honestly significant differences (HSD) procedure was used. To aid the interpretation of interactions, we also performed contrast-analyses for sub-interactions where it provided information that was complementary to traditional pairwise post-hoc tests. Statistical analysis was done using STATISTICA 13.1 (StatSoft, Tulsa, OK).

Peak-based statistics might be more tolerant to latency differences of well-defined sharp peaks, but it is known that they do not provide a comprehensive view on the temporal evolution of an effect (Rousselet & Pernet, 2011), also, peaks do not necessarily coincide with the latent components that ERP research aims to uncover (Kappenman & Luck, 2011; Luck, 2005). On the individual ERP curves, particularly in peripheral stimulation conditions, P1 peaks were frequently ambiguously definable, probably due to the well-known variability of these early components, related to cortical folding (Ales, Carney, & Klein, 2010; Ales, Yates, & Norcia, 2010). Also, even for central stimulation when relatively clear P1 peaks were found, inspecting the difference curves of the inversion effect revealed a mismatch between the latency where the effect is strongest and the peak latencies of the grand averages (Grand average peak latencies on the P1 electrode cluster were the following: left, upright: 110 ms; left, inverted: 118 ms; right, upright: 107 ms; right, inverted: 115 ms; Median latencies of individually detected peaks were the following: left, upright: 109 ms; left, inverted: 120 ms; right, upright: 106 ms; right, inverted: 111 ms; see also Figure 5.C). In particular, inversion rather seemed to influence the descending edge after the P1 peak towards the N170 (peak of the difference curve of the central inverson effect on the pooled P1-N170 electrode cluster, left: 140 ms; right: 135 ms; see Fig. 5C).

For these reasons we also conducted a mass univariate time-series analysis to uncover further possible patterns in the inversion effect that could be obscured by the weaknesses of the peak-based approach. This analysis was performed separately for central and peripheral stimulation conditions, and focused on inversion effects and their interactions with hemispheric and visual field laterality in the first 250 ms of ERP on the bilateral pooled P1-N170 electrode clusters, i.e. the union of the electrode clusters of the P1 and the N170 component defined above (P7, P9, PO7, PO9, O1, and O9 for the left and P8, P10, PO8, PO10, O2, and O10). For each test, the type I error inflation caused by testing at each time point was corrected for by means of the cluster-based permutation method as implemented in the FieldTrip toolbox (Maris & Oostenveld, 2007; Oostenveld, Fries, Maris, & Schoffelen, 2011). The cluster threshold was set at the p=0.05 two-tailed significance level of the t-test conducted at each time point (df=15). The maximum-sum cluster statistic was used, and 9999 permutations were performed. The resulting p values, reported as p_Cluster_, were multiplied by 2 to account for two-tailed permutation testing.

#### 2.4.4. Eye Tracking Data Acquisition and Analysis

Eye movements were recorded at a 1250 Hz sampling rate using iViewX Hi-Speed tracking column (SMI GmbH, Teltow, Germany) during the EEG experiment. Eye tracking data was segmented into 500 ms epochs starting 200 ms preceding stimulus onset. We applied baseline correction by subtracting the average gaze position in the 100 ms preceding stimulus onset, assuming that participants fixated the fixation dot before the appearance of the face stimulus (Knakker, Weiss, & Vidnyánszky, 2015). Subsequently, we calculated an index to identify participants with too much eye movement. We used the common interval of the N170 component (150-200 ms after stimulus onset) assuming that eye movements in this period would lead to ocular EEG artefacts affecting the N170 ERP component. We defined the threshold of the shift from the fixation dot in 1.5° (half of the distance from the fixation point to the edge of the central stimulus) for each time point. If a trial in this interval contained any time point more than 1.5° away from the fixation dot, it was marked as incorrect fixation. If more than 90% of trials were marked because of inaccurate fixation, the participant was excluded from all further analyses (n=2).

### 2.5. Foveal masking experiment

#### 2.5.1. Participants

Ten healthy participants attended the experiment (1 male, 1 left-handed, mean ± SD age: 23.18 ± 2.4 years). One participant completed the first three blocks only. Each subject had normal or corrected-to-normal vision. None of them reported any history of neurological or psychiatric disease. They provided written informed consent in accordance with the protocols approved by the United Ethical Review Committee for Research in Psychology (EPKEB), Budapest, Hungary.

#### 2.5.2. Stimuli and Procedure

Participants performed the same 3AFC task as described above, but a dynamic mask was presented at the center of the screen. The masks were upright faces, inverted faces or noise patches. In one fourth of the trials, no mask was shown. Five new face identities were generated using FaceGen, while noise patches were created by phase-scrabling with the minimum-phase method as defined by Ales et al. (Ales, Farzin, Rossion, & Norcia, 2012). Faces were unmorphed to obtain better performance for peripheral stimuli than in the first experiment. Target faces were presented only on the periphery (LVF, RVF), scaled according to the CMF the same way as in the main experiment (11.17° × 14.9°), while the dynamic mask, placed in the center was shown at the same size as the central stimuli (3° × 4°) of the former experiment (Fig.1.).

The training phase was also similar to the one during the EEG experiment. First, the three identities were shown side by side to allow familiarization with the target stimuli. Then, a practice phase was run as described for the main experiment, with the target faces presented individually in the center of the screen. This phase was followed by an extra practice phase to get participants acquainted with the fast peripheral stimulus presentation as well: stimulus presentation was shorter than in the main experiment (100 ms) and they appeared only on the periphery. No mask was presented during this training phase. After reaching a performance higher than 75%, participants commenced to the test phase.

During the test phase, each trial started with a fixation point in the center of the screen, which was presented through the entire trial. The target stimulus appeared for 100 ms and was followed by a dynamic mask centered at fixation at one of five possible SOAs (50 ms, 100 ms, 150 ms, 200 ms, or 250 ms). The mask was presented for 83 ms (5 frames), during which in every odd frame a new identity or noise patch was shown. Participants had to indicate face identity by pressing one of three keys (‘left arrow’, ‘bottom arrow’ or ‘right arrow’ on a QWERTY keyboard) as fast and accurately as possible. They had 2000 ms to make their response after stimulus offset. Participants received feedback by the fixation point changing its color. The behavioral experiment consisted of 4 blocks with short breaks in the middle of blocks. Altogether, the test consisted of 1920 trials leading to 24 trials per condition. Participants viewed the stimuli on a 24 inch LCD monitor at a refresh rate of 60 Hz from 66 cm. The experiment lasted approximately two and a half hours.

#### 2.5.3. Analysis of the Foveal Masking Experiment

Similarly to the EEG experiment, the responses and reaction times (RTs) were collected and were submitted to a repeated-measures analysis of variance (ANOVA) with Orientation (UP, INV), Position (LVF, RVF), Mask type (Noise, Upright and Inverted face, No mask) and SOA (5 levels: 50 ms, 100 ms, 150 ms, 200 ms, 250 ms) as within-subject factors. The Greenhouse-Geisser correction was applied to correct for possible violations of sphericity. Post hoc tests were computed using Tukey’s honestly significant differences (HSD) procedure. Statistical analysis was carried out using STATISTICA 13.1 (StatSoft, Tulsa, OK).

Eye tracking data acquisition and analysis was very similar to the acquisition and analysis of the EEG experiment, except that the eye movements were recorded at a 240 Hz sampling rate using a different iViewX Hi-Speed tracking column. Preprocessing of the acquired data was done as described above for the main experiment. None of the participants had to be excluded because of inadequate fixation. Further analysis was conducted identically to the previous experiment, and mean fixation stability values between 0-100 ms for each condition were entered into a two-way repeated-measure ANOVA with stimulus Orientation (UP, INV) and Position (LVF, RVF) as within-subject factors.

To assess the stability of fixation during stimulus presentation and validate that it did not differ significantly between conditions, mean distance from fixation was calculated across trials for all time points and averaged from onset to offset of the stimulus (0-150 ms) for each condition. Mean fixation stability values were analyzed with repeated-measures analysis of variance (ANOVA) with stimulus Orientation (UP, INV) and Position (LVF, C, RVF) as within-subject factors. Statistical analysis was done using STATISTICA 13.1 (StatSoft, StatSoft, Tulsa, OK).

## 3. Results

### 3.1. Behavioral results

The behavioral results revealed (Fig. 2A) that face identity discrimination performance was significantly better for upright than for inverted faces (main effect of Orientation: F(1,15)=108.66, p<0.001, η^2^=0.88). Importantly, the strength of this FIE was similar for foveal and peripheral faces (Orientation and Position interaction: F(1.63,24.41)=0.24, p=0.75, η^2^=0.81), even though the overall performance was lower at the periphery (main effect of Position: F(1.34,20.10)=85.78, p<0.001, η^2^=0.67; post hoc tests for periphery (LVF, RVF) vs fovea all p<0.001). No differences were found in the discrimination accuracy for peripheral faces between the left and right visual fields (post hoc test for LVF vs RVF: p=0.73). Face inversion also had an effect on the participants’ reaction times (main effect of Inversion: F(1,15)=6.78, p=0.020, η^2^=0.31). However, participants made faster responses for upright than inverted faces only when the stimuli were presentted foveally, but not in the case of peripheral faces (Fig. 2B, Orientation and Position interaction: F(1.79,26.85)=7.28, p=0.004, η^2^=0.89, post hoc test for C: p<0.001, in the case of periphery all p>0.05).

**Fig.2.**
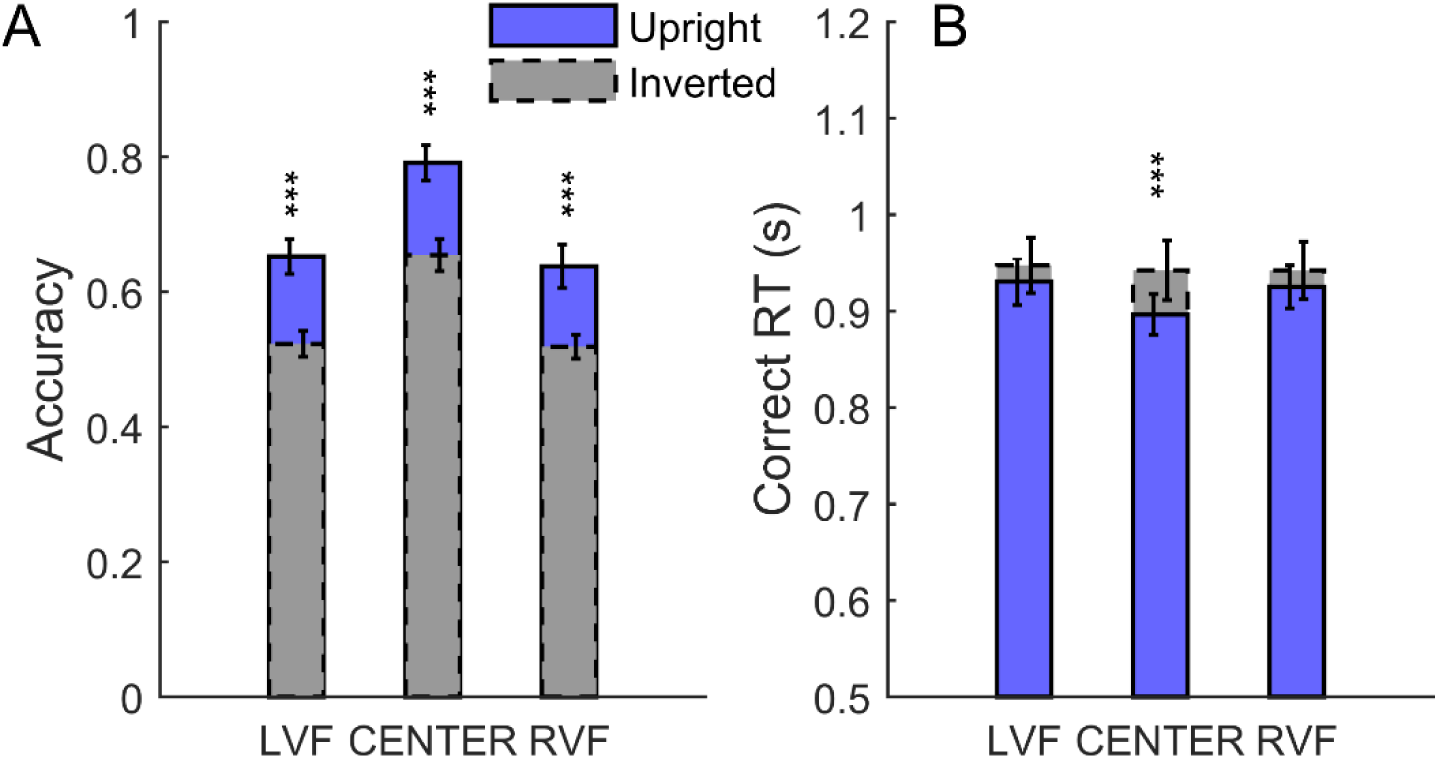
Accuracy (A) and correct reaction times (B) of the three-alternative forced-choice identity-discrimination task during EEG. A. Behavioral FIE was present independently of position. B. Responses were significantly faster for upright faces when the stimulus was presented in the center. Error bars indicate ± SEM (N=16 in both cases, ***p<0.001).

Next, we performed a correlation analysis to explore the relationship between foveal and peripheral FIEs. The results of this analysis revealed a strong positive correlation in the strength of the FIE (difference in accuracy between upright and inverted faces) between the peripheral face stimuli presented in the left and right hemisphere (Fig.3.; LVF vs RVF: r_(14)_=0.60, p=0.014, CI=[0.35 0.81]; NO=0). On the other hand, no significant correlation was found in the strength of FIE between the foveally and peripherally presented faces (Fig.3.; LVF ~ C: r_(14)_= −0.22, p=0.41, CI=[-0.53 0.14], NO=0; RVF ~ C: r_(14)_= −0.29, p=0.28, CI=[−0.61 0.28], NO=0). We also compared the correlation coefficients using Zou’s confidence interval (Zou, 2007) and found that the correlation in FIEs between the LVF and RVF faces differs significantly from the correlations between the foveal and peripheral face stimuli (LVF ~ C vs LVF ~ RVF: CI=[0.15 1.38]; RVF ~ C vs LVF ~ RVF: CI=[0.05 1.36]). Taken together, the behavioral results suggest that peripheral face perception involves holistic processing of facial attributes, which might differ from those involved in the perception of foveal faces.

**Fig.3.**
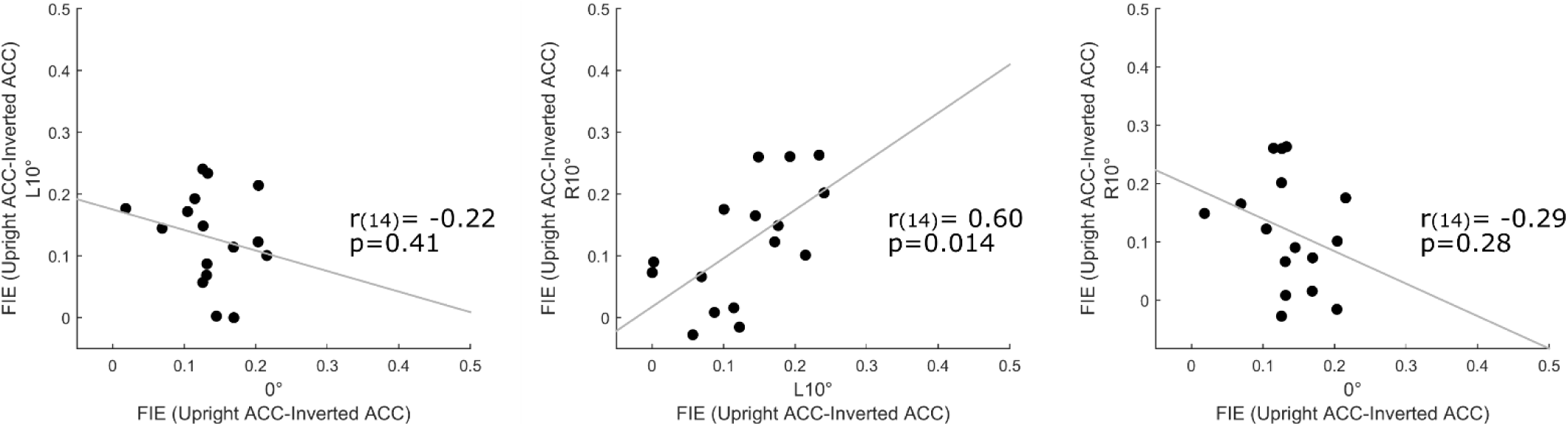
Relationship between behavioral face inversion effects at each position. Dots represent individual subjects (N=16). The left and the right panels show the relationship between foveal and peripheral FIEs, in LVF and RVF, respectively. The middle panel shows significant correlation between the FIE of the LVF and the RVF. Diagonal lines indicate linear least-squares fit.

### 3.2. ERP results

First, we performed an analysis of the EEG data to reveal the effect of face inversion on the early P1 and N170 ERP components evoked by foveal and peripheral faces (Fig. 4.).

**Fig.4.**
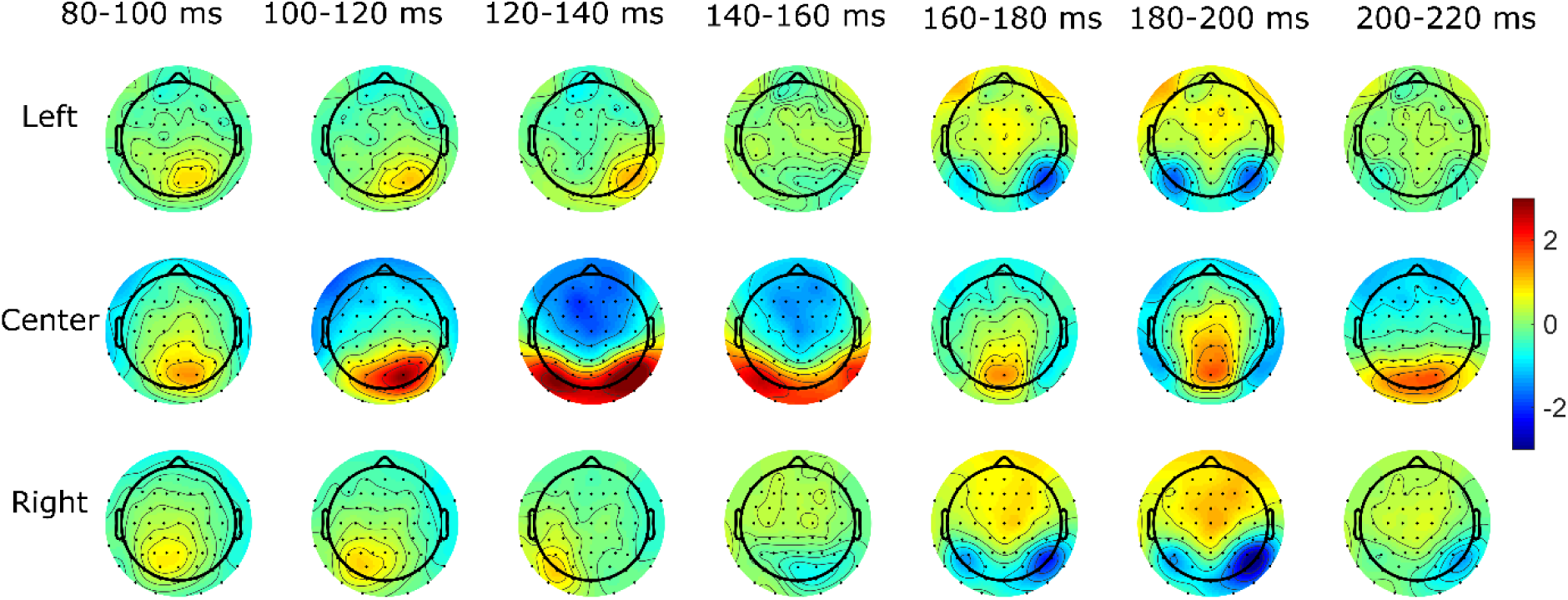
The temporal evolution of the scalp difference topographies (inverted-upright) during the time course of the P1 and N170 component.

The results revealed strong, but markedly different face inversion effects on the amplitude of the P1 and N170 components (Fig. 5.B). P1 amplitudes were increased for inverted as compared to upright faces only for the foveally presented faces (Orientation × Position interaction F(1.11,16.60)=7.08, p=0.015, η^2^=0.55; post hoc tests for upright vs inverted faces presented ipsilaterally: p=0.99, centrally: p<0.001, contralaterally: p=0.89). Our finding showing that the FIE on the P1 component differs between foveal and peripheral faces was also confirmed by contrast analysis (Orientation effect on periphery vs center: F(1,15)=7.41, p=0.016). On the other hand, we found that FIE on the N170 amplitudes was significant for both foveal and peripheral faces, with the peripheral FIE being stronger than the foveal (main effect of Orientation: F(1,15)=21.17, p<0.001, η^2^=0.59; Orientation × Position interaction: F(1.25,18.82)=5.22, p=0.028, η^2^=0.63; post hoc tests: upright vs inverted for faces presented ipsilaterally: p<0.001, for centrally: p=0.041, for contralaterally: p<0.001). Although there was a significant FIE on the N170 for all three positions, we tested whether the magnitude of the FIEs differ among positions. Contrast analysis revealed that FIE between ipsilateral and foveal presentation was different (F(1,15)=6.72, p=0.020) with greater FIE for ipsilaterally presented stimuli, while contralateral and foveal FIE did not differ from each other (F(1,15)=1.29, p=0.27). Ipsilateral FIE was greater than FIE for contralateral stimulus (F(1,15)= 15.22, p=0.001). Furthermore, in line with previous research showing a right hemisphere lateralization of visual cortical face processing, we found stronger face inversion effects on N170 amplitudes over the right as compared to the left hemisphere (Hemisphere × Orientation interaction: F(1,15)=10.19, p=0.006, η^2^=0.40, post hoc tests for upright vs inverted for LH: p<0.001; for RH: p<0.001).

**Fig.5.**
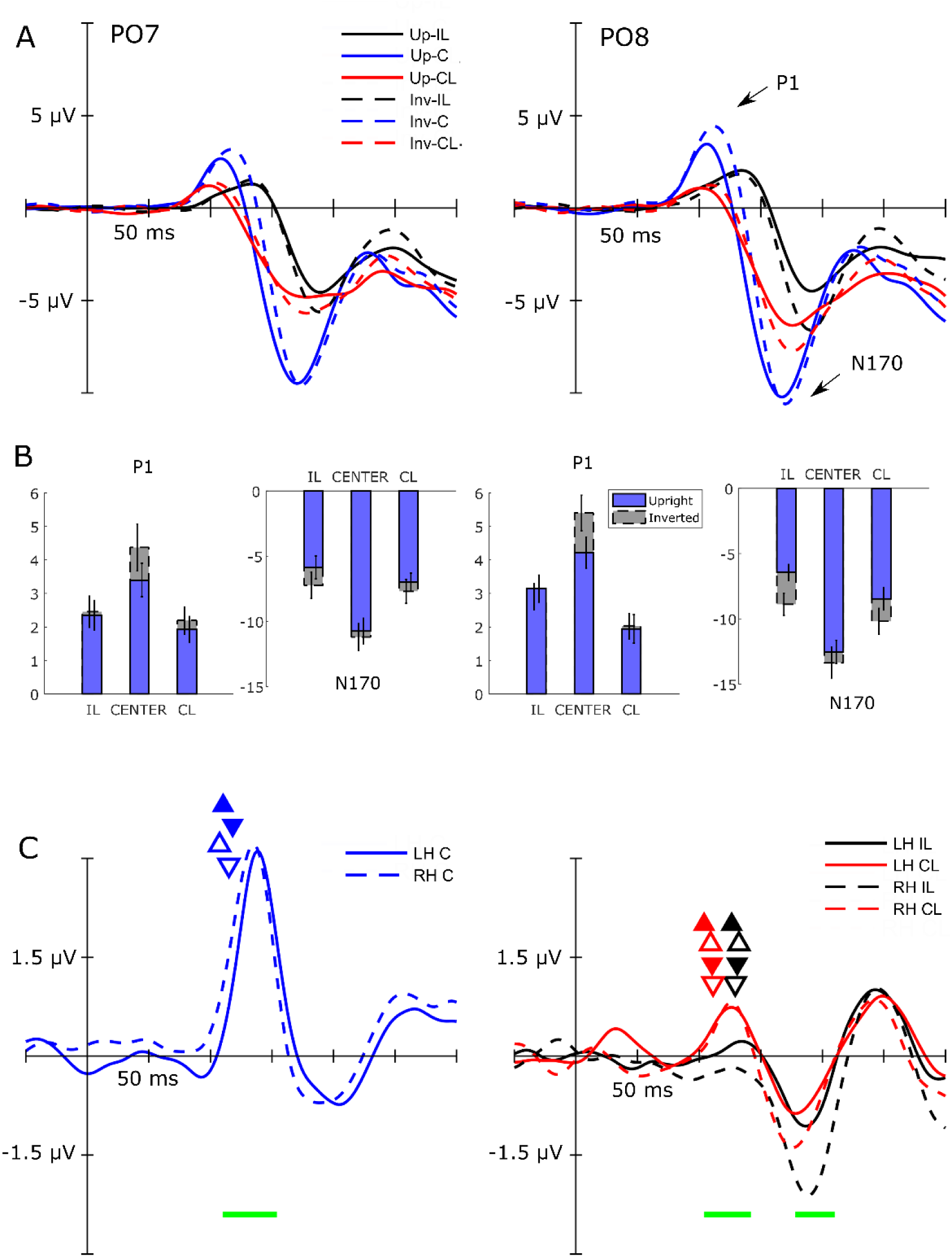
**A.** Grand average ERP waveforms at two occipital electodes (PO7, left and PO8 right hemisphere respectively). ERPs are shown as a function of Position (IL, CL, C) and Orientation (UP, INV) **B**. Results of the ERP analysis. Blue bars indicitace amplitude for upright faces, while grey bars show amplitude of ERP responses in the case of inverted faces. Results are derived from the clusters of electrodes (see Methods). Bars are arranged according to the hemispheres and components (LH and RH, P1 and N170 respectively). Error bars indicate ± SEM (N=16 in both cases, *p<0.05, ***p<0.001). **C.** ERP difference waveforms at the union of the electrode clusters of the P1 and the N170 component plotted according to positions. On the left panel, ERPs elicited by inverted faces minus the ERPs of upright faces for central position are presented. On the right panel, difference waves are presented as a function of peripheral presentation averaged across electrodes. Horizontal green lines under the difference curves represent the time ranges during which the time point by time point analysis showed significant inversion effects (110-154 ms, 105-140 ms and 180-210 ms respectively). Up-pointing triangles indicate the mean peak latency for upright faces, while down-pointing triangles show the mean P1 latency for inverted faces. Filled triangles indicate the values on the left hemisphere, empty triangles show latency values on the right hemisphere.

To reveal the temporal evolution of the FIE, we also conducted a point-by-point analysis on the difference curves of the inversion effect. This was important because the visual inspection of the timecourse of FIE on the scalp topoghraphy maps suggested that the time window of the strongest FIE does not match with the grand average peaks of the P1 component (for observed discrepancies see Methods section and Fig. 5C.).

Indeed, for centrally presented face stimuli, the timepoint-by-timepoint analysis (see Methods section) revealed a significant FIE in a time interval ranging between 110-155 ms following the stimulus onset and corresponding to the descending edge of the P1 component (INV > UP, 110-155 ms, p_Cluster_=0.0004, Fig. 5C.). Importantly, on the periphery, a similar effect was only present for contralaterally, but not for ipsilaterally presented stimuli (IL vs CL × Orientation interaction: 105-140 ms, p_Cluster_=0.002 see Fig.5C; simple inversion effect for CL stimuli: 110-140 ms, p_Cluster_=0.04; no suprathreshold timepoints for IL stimuli).

The inversion effects in the time window of the N170 component were detected using this method only for peripheral (CL: 155-195 ms, p_Cluster_=0.007, IL: 155-205 ms, p_Cluster_=0.005), but not for centrally presented stimuli (p_Cluster_>0.05), probably due to the different sensitivity profile of the peak-based and the cluster-based methods (see Methods section). The timepoint-by-time point analysis also suggested an ipsilateral bias of the N170 inversion effects (IL vs CL × Orientation interaction: 180-210 ms, p_Cluster_=0.017; Fig.5C). The right-hemisphere lateralization of the N170 inversion effects found in the peak-based analysis was also present (Hemisphere × Orientation interaction: 140-200 ms, p_Cluster_=0.012).

In sum, the electrophysiological results are in close agreement with our behavioral findings. First, similarly to discrimination performance, early ERP responses were affected by face inversion for both foveal and peripheral faces. Second, the possibility that foveal and peripheral inversion effects might be mediated by different processes – suggested by the results of the correlation analysis of the behavioral data – received further support from the ERP results showing that the effect of face inversion differs for foveal and peripheral faces, the former being dominant in the P1 while the latter more in the N170 time-windows.

### 3.3. Results of the foveal masking experiment

In accordance with the behavioral results of the EEG experiment, we found strong FIEs for peripheral stimuli: face identity discrimination performance was significantly better for upright than for inverted faces (main effect of Orientation: F(1,9)=85.71, p<0.001, η^2^=0.90). It was also found that participants’ performance differed across the different mask types used during the task (main effect of Mask type: F(1.8,16.21)=17.44, p<0.001, η^2^=0.60). Interestingly, visual noise masks had no effect on discrimination performance (post hoc p=0.69), however, applying upright and inverted face masks disrupted performance significantly when compared to the no mask condition (post hoc tests for both face masks, p<0.001). Performance between the upright and inverted face mask conditions did not differ significantly (F(1,9)=2.89, p=0.12). Our results also revealed that performance differed across SOAs (F(2.55,22.98)=3.78, p=0.030, η^2^=0.64), performance was better when the mask appeared with 250 ms SOA compared to 50 ms and 100 ms (post hoc, 250 ms vs 50 ms SOA: p=0.031; 250 ms vs 100 ms SOA: p=0.049). Most importantly, however, the magnitude of the peripheral FIE was not affected by foveal face mask stimuli (Orientation × Mask interaction: F(1.95,17.52)=1.55, p=0.24, η^2^=0.65; see Fig. 6.).

**Fig.6.**
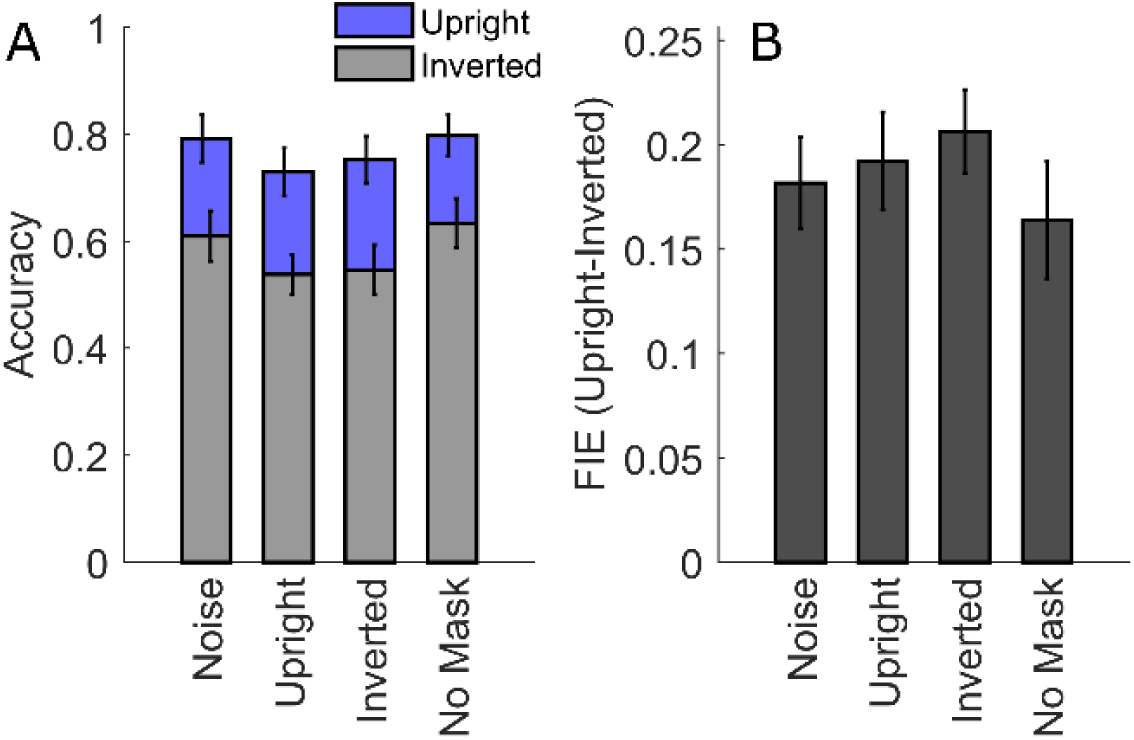
Accuracy (A) and FIE indices (B) from the foveal masking experiment. Blue bars indicate accuracy for upright target faces, while grey bars show accuracy for inverted target faces for the different mask conditions, separately. Error bars indicate ± SEM (N=10 in both cases).

### 3.4. Results of the eye movement data analysis

We measured participants’ gaze position to assess whether they maintained fixation at the center of the screen during both the EEG and foveal masking experiments. During the EEG experiment, all participants fixated properly at least in 97% of the trials during stimulus presentation (0-150 ms). Mean distance from the fixation dot was 0.27°, and it did not differ significantly between conditions (main effect of Orientation: F(1,15)=0.193, p=0.67; main effect of Position: F(1.54, 23.17)= 2.43, p=0.12, η^2^=0.77; interaction of Orientation and Position: F(1.99,29.81)= 0.03, p=0.97, η^2^=0.99).

Gaze position was monitored during the foveal masking experiment as well. We analyzed whether they fixated on the center of the screen during the experiment independently of the presentation site. All participants fixated properly at the stimulus at least in 94% of the trials during stimulus presentation (0-100 ms). Mean distance from the fixation dot was 0.22°, and it did not differ significantly between conditions (main effect of Orientation: F(1,6)=0.22, p=0.66, η^2^=0.58; main effect of Position: F(1,6)= 1.98, p=0.21, η^2^=0.25; main effect of mask: F(1.36,8.13)= 0.89, p=0.41, η^2^=0.45, main effect of SOA: F(2.02,12.13)=1.59, p=0.24, η^2^=0.51).

## 4. Discussion

The results revealed strong face inversion effects on the behavioral as well as on the ERP responses when faces were presented in the periphery. Discrimination of the identity of inverted peripheral faces was strongly impaired as compared to upright faces, and the magnitude of this impairment was similar for centrally and peripherally presented faces. The early ERP responses were affected by inversion in two time intervals. The first component of the FIE was peaking after the P1 component, between 130-140 ms following stimulus presentation for both centrally and peripherally displayed faces, and in the latter case it was present only over the contralateral hemisphere. The timing of the second FIE component corresponded closely with the time window of the N170 component and it showed a right hemisphere lateralization for peripheral faces. Furthermore, our foveal masking experiment revealed that upright and inverted face masks strongly impair peripheral face identity discrimination performance, but do not reduce the magnitude of the FIE.

The face inversion effect is one of the most widely used markers of expert face-selective processing, and the extensive behavioral research suggests that impairment of face perception in the case of inverted faces might be primarily due to the disruption of holistic face processing enabling the integration of the facial features into a whole face (Piepers & Robbins, 2012; Rossion, 2008, 2009)^1^. Previous EEG research showed that face inversion affects the early components of the ERP responses, including the P1 as well as the N170 component, associated with structural processing of facial information (Bentin et al., 1996; Eimer, 2000; Jacques et al., 2007; Rossion et al., 2000). Both P1 and N170 elicited by inverted faces emerge later with increased amplitudes relative to upright faces (Bentin et al., 1996; Eimer, 2000; Jacques et al., 2007; Linkenkaer-Hansen et al., 1998; Rossion et al., 2000). Furthermore, functional magnetic resonance imaging (fMRI) research provided evidence for face inversion effects within the visual cortical regions of the core face network, such as the face-selective FFA (for review see Yovel, 2016), as well as in regions outside the core network (Matsuyoshi et al., 2015; Rosenthal, Sporns, & Avidan, 2016; Zachariou, Nikas, Safiullah, Gotts, & Ungerleider, 2016), including the object-selective lateral occipital cortex (LO) (Grill-Spector, Kushnir, Edelman, Itzchak, & Malach, 1998; Larsson & Heeger, 2006) and the intraparietal sulcus (Haxby et al., 2000; Haxby, Hoffman, & Gobbini, 2002; E. A. Hoffman & Haxby, 2000). The involvement of LO in the processing of inverted faces was also strongly supported by a recent study (Rosburg et al., 2010) using intracranial electrophysiological recordings and showing increased neural response amplitudes in the LO for inverted faces compared to upright faces. Taken together, these findings are consistent with the view that the processing of upright and upside down faces differs both quantitatively and qualitatively. According to this account, inversion effects on the fMRI responses within the face network might reflect the failure of face-selective holistic processing of inverted faces, resulting in increased demands for sequential processing of facial features and the extraction of relational information among them (Goffaux & Rossion, 2007; Rossion, 2008). This, in turn, will be reflected in the modulation of the neural responses within the face network as well as in the engagement of object-selective visual cortical processes within the LO (Matsuyoshi et al., 2015; Rosburg et al., 2010) and visuospatial mechanisms of the dorsal visual stream and the parietal cortex (Zachariou et al., 2016). Based on these experimental findings and theoretical considerations it has been proposed that face inversion effects on the EEG/MEG and fMRI responses might be used as neural markers of holistic face processing (for review see Yovel, 2016).

Up to now, research on face perception focused primarily on the processing of faces presented in the central visual field. This might not be surprising, considering previous suggestions that visual object processing in the human ventral temporal cortex (VTC) is spatiotopically organized and representation of objects whose recognition depends on the analysis of fine details - such as faces - is associated with a central visual-field bias (Amedi et al., 2001; Grill-Spector & Weiner, 2014; Hasson et al., 2002; Kanwisher, 2001; Levy, Hasson, Avidan, Hendler, & Malach, 2001; Malach et al., 2002). It has been shown that in FFA nearly all neural resources are dedicated to the central (~7 deg) portion of the visual field (Kay et al., 2015). Therefore, it may seem justified to neglect the question of holistic processing on peripheral faces. However, our everyday social interactions very often involve situations when more than two persons are interacting simultaneously. In this case, one face is fixated at a time, whereas the remaining faces are located in the peripheral visual field. Importantly, not even these peripheral faces can be neglected, since they may carry important, socially relevant information, such as changes in facial expression. This, on the other hand, implies that humans are highly experienced in peripheral face perception and raises the possibility that holistic processing might also be involved in the perception of peripheral faces.

Indeed, the possibility that peripheral face perception might also involve face-selective neural processes received support from a previous study showing that larger N170 amplitudes for faces than houses - a marker of face-selective processing - can be observed for peripheral faces if the size of the stimuli is scaled according to the V1 cortical magnification factor (Rousselet, Husk, Bennett, & Sekuler, 2005). Furthermore, the involvement of holistic processing for peripheral faces has been suggested by a previous behavioral study, showing a strong face inversion effect for peripheral faces (McKone, 2004). Importantly, in a control experiment it was also found that inversion had no effect on the identification of an isolated face feature (nose), suggesting that peripheral FIE, similarly to that found in the central visual field, might indeed result from the disruption of holistic face perception (McKone, 2004). In accordance with these previous findings, the present study revealed that perception of inverted peripheral faces is impaired as compared to upright faces. We also found a significant correlation between the magnitudes of FIEs from the left and right peripheral visual fields, whereas no association was found between the size of peripheral and central FIE. This suggests that the neural substrate of the central and peripheral FIE might be different. Furthermore, our ERP findings provide the first evidence that face inversion affects early neural responses to peripheral faces. Neural responses for upright and inverted peripheral faces differed in two temporal intervals, between 110-150 ms and 160-210 ms following the stimulus onset. The two peripheral FIE components differed in their scalp distribution: the earlier component was present only over the contralateral hemisphere, whereas the later showed strong right hemisphere lateralization, independently of whether faces were displayed in the left or right visual field. These results fit well into the interpretation framework of face inversion effects in the central visual field, described above. Although previous research investigating inversion effects on the P1 ERP component for centrally presented faces obtained mixed results (for review see Yovel, 2016), a very recent study showed a robust FIE for central faces in a time window (peaking between 130-140 ms following the stimulus onset) that is in a remarkable agreement with the temporal interval of the earlier FIE component found in the present study for both central and peripheral faces (Colombatto & McCarthy, 2016). In both this recent study and our work, P1 peaks and the maximum of the early central inversion effect were clearly dissociated, which, together with findings of similar peak-effect discrepancies (Rousselet, Macé, Thorpe, & Fabre-Thorpe, 2007) and theoretical considerations (Kappenman & Luck, 2011; Luck, 2005; Rousselet & Pernet, 2011), highlight the importance of going behind peak-based analyses to improve the sensitivity of ERP experiments. Our findings showing that this earlier FIE component is confined to the contralateral hemisphere for peripheral faces imply that the source of this component might be localized in retinotopically organized visual cortical regions with neural receptive fields representing the contralateral visual field (Towler & Eimer, 2015). Since it has been shown that LO consists retinotopically organized subregions (Larsson & Heeger, 2006), it is tempting to assume that the early peripheral FIE component found in the present study might signal the proposed qualitative difference in the processing of inverted and upright faces; i.e., the involvement of the object-selective neural processes of the LO in the processing of inverted faces.

The timing of the later peripheral FIE component closely corresponded to the time window of the N170 component. Previous studies have consistently reported increased N170 amplitudes and latencies for inverted as compared to upright faces presented centrally (Bentin et al., 1996; Eimer, 2000; Jacques et al., 2007; Rossion et al., 2000; Rossion et al., 2003) and our results for centrally presented faces provided yet another example of supporting evidence. Although it has been shown that simultaneous neural activity within several visual cortical regions (including the FFA, OFA and STS) might contribute to the N170 component (Rossion, 2014; Rossion & Jacques, 2008), the most commonly held view is that inversion effects on the N170 component originate from the regions of the core face network and thus might reflect increased processing demands required for the integration of relational information among facial features (Goffaux & Rossion, 2007; Rossion, 2008) in the case of inverted faces as compared to upright faces (quantitative difference). This interpretation of the peripheral FIE found in the present study is supported by its scalp distribution. In contrast to the earlier FIE component, which was present only contralaterally, the later peripheral FIE component showed a strong right hemisphere lateralization, independently of whether the faces were displayed in the left or in the right visual fields. This is in agreement with the right hemisphere dominance in face-selective visual cortical processing (Jacques & Rossion, 2009; Pitcher, Walsh, Yovel, & Duchaine, 2007; Rossion et al., 2000; for review see Yovel, 2016) and suggests that the later peripheral FIE component might reflect increased processing demands within the face network required for the integration of relational information among facial features (Goffaux & Rossion, 2007; Rossion, 2008).

Traditionally, it is believed that the coding of peripheral objects starts at the lower-level retinotopically organized visual cortical regions and proceeds towards higher-level, object-selective VTC areas with more position-tolerant object representations (DiCarlo & Cox, 2007; Hoffman & Logothetis, 2009; Riesenhuber & Poggio, 2000; Rousselet, Thorpe, & Fabre-Thorpe, 2004). However, recent research provided evidence suggesting that peripheral object perception might involve a foveal processing component that is mediated by feedback signals to the visual cortex representing the central visual field, which can be revealed by presenting foveal noise shortly after the presentation of the peripheral target objects (Fan et al., 2016; Williams et al., 2008). Therefore, we performed a behavioral experiment to test the possibility that neural processes dedicated to the central visual field are involved in the face inversion effect obtained for peripheral face stimuli. We reasoned that if peripheral holistic face processing is mediated by neural processes dedicated to the central visual field, adding foveal masks, in addition to modulating the overall face discrimination performance, should strongly reduce the size of peripheral FIE. Our results revealed that foveal upright and inverted face masks, but not noise masks impair peripheral identity discrimination performance strongly for both upright and inverted faces. Importantly, however, the magnitude of the peripheral FIE was not affected by the foveal face masks. These results suggest that holistic processing of peripheral faces is not compromised when in the same time face-selective neural resources dedicated to the central visual field are engaged in processing of an irrelevant foveal face mask, suggesting that it might be mediated by neural processes within the visual cortex representing the peripheral visual field.

An important question is how the results of the present study can be reconciled with the previous neuroimaging findings (Amedi, Malach, Hendler, Peled, & Zohary, 2001; Grill-Spector & Weiner, 2014; Hasson, Levy, Behrmann, Hendler, & Malach, 2002) showing that in the face-selective regions of the ventral temporal cortex, neural resources involved in holistic processing are primarily dedicated to the central portion of the visual field. There is an important difference in the stimulus design between these previous and the current study that might help to explain the discrepancies in the obtained results. In our study, peripherally presented faces were scaled according to the cortical magnification factor, whereas in the previous studies showing central visual field dominance, no such scaling was applied. It has been shown that both the magnitude of the fMRI responses in FFA (Yue, Cassidy, Devaney, Holt, & Tootell, 2011) and the amplitude of the N170 component (Rousselet et al., 2005) are affected by the size of peripheral face stimuli, and that only in the case when peripheral stimuli are scaled according to the cortical magnification will these face-selective neural markers show similar results for peripherally and centrally presented faces. Therefore, using unscaled peripheral stimuli might be one of the reasons why in previous neuroimaging research face-selective representations in VTC were found to be much weaker in the peripheral as compared to central visual field.

To conclude, our findings suggest that faces are processed holistically throughout the visual field, and pave the way for further research aiming at uncovering the similarities and differences in the neural processes underlying central and peripheral holistic face processing.

## Acknowledgement

This work was supported by a grant from the Hungarian Brain Research Program (KTIA_13_NAP–A–I/18) to Zoltan Vidnyánszky.

## Footnotes

1 It must be noted that facial feature discrimination performance is also deteriorated in case of inverted as compared to upright faces (ref)

